# Spillover of H5 influenza viruses to vampire bats at the marine-terrestrial interface

**DOI:** 10.1101/2025.11.09.686930

**Authors:** I-Ting Tu, Christina Lynggaard, Lorin Adams, Sarah K Walsh, Hanting Chen, Savitha Raveendran, Matthew L Turnbull, Megan E Griffiths, Rita Ribeiro, Jocelyn G Peréz, William Valderrama Bazan, Carlos Tello, Carlos Zariquiey, Kristhie Pillaca Rodriguez, Marco Risco, Illariy Quintero Mamani, Wendi Chávez, Roselvira Zuniga Villafuerte, Joaquin Clavijo Manuttupa, Jean Pierre Castro Namuche, Andres Moreira Soto, Jan Felix Drexler, Gustavo Delhon, Christina Faust, Susana Cárdenas-Alayza, Ed Hutchinson, Pablo R Murcia, Massimo Palmarini, Kristine Bohmann, Ruth Harvey, Daniel G Streicker

**Affiliations:** School of Biodiversity, One Health and Veterinary Medicine, University of Glasgow, Glasgow, UK; MRC-University of Glasgow Centre for Virus Research, Glasgow, UK; Globe Institute, Faculty of Health and Medical Sciences, University of Copenhagen, Copenhagen, Denmark; Worldwide Influenza Centre, The Francis Crick Institute, London, UK; Royal Veterinary College, London, UK; Universidad Peruana Cayetano Heredia. Facultad de Medicina Veterinaria y Zootecnia, Lima, Perú; Asociación para el Desarrollo y Conservación de los Recursos Naturales (ILLARIY), Lima, Perú; Charité Universitätsmedizin, Berlin, Germany; University of Nebraska-Lincoln, NE, USA; Centro para la Sostenibilidad Ambiental, Universidad Peruana Cayetano Heredia, Lima, Perú; Facultad de Ciencias e Ingeniería, Universidad Peruana Cayetano Heredia, Lima, Perú; Programa de Maestría en Salud Pública y Salud Global. Facultad de Salud Pública y Administración. Universidad Peruana Cayetano Heredia, Lima, Perú; Centro de ornitología y biodiversidad (CORBIDI), División de Mastozoología, Lima, Perú; Department of Viroscience, Erasmus Medical Center, Rotterdam, The Netherlands

## Abstract

The highly pathogenic H5N1 avian influenza A virus (IAV) clade 2.3.4.4b has spread globally and spilled over into multiple mammalian species, raising concerns about its pandemic potential. In late 2022, clade 2.3.4.4b viruses devastated seabird and marine mammal populations along the Pacific coast of South America. Here, we report the first evidence of H5 IAV infections in wild bats globally, focusing on common vampire bats (*Desmodus rotundus*) in coastal areas of Peru. Longitudinal serological screening, stable isotope analysis and metabarcoding revealed repeated exposures to H5 IAVs in vampire bats which feed on coastal wildlife species heavily impacted by the 2.3.4.4b epizootic, but no evidence of infection in populations without access to marine prey. We further report bat gene flow between IAV-exposed and IAV-naïve populations, and IAV infections in a vampire bat colony that fed on both marine and terrestrial livestock prey, providing insights into how future IAV epizootics might spread spatially within bats and between marine and terrestrial ecosystems if a bat reservoir were established. Immunohistochemistry demonstrated that the H5 haemagglutinin protein binds to the upper respiratory tract of vampire bats, suggesting bat tissue susceptibility to H5 IAVs. Finally, vampire bat-derived kidney, liver, and lung cells supported entry, replication, and egress of avian and mammalian 2.3.4.4b viruses, confirming cellular infectivity. These results illustrate how combining ecological inference and experimental virology can pinpoint the species origins and biological significance of viral spillover at species interfaces. Recurrent exposures from marine wildlife, tissue and cellular susceptibility to H5N1 IAVs, and connections to other IAV-susceptible terrestrial mammals establish the prerequisite conditions for vampire bats to spread IAVs between marine and terrestrial environments or to form a novel reservoir of highly pathogenic IAVs.

## Main text

Highly pathogenic avian influenza A viruses (H5N1) clade 2.3.4.4b have raised pandemic concerns due to their rapid intercontinental spread and transmission in mammals, including dairy cattle in the United States^1^ and South American sea lions (*Otaria byronia*)^2^. As of 10 November 2025, 70 human cases of bovine-origin H5N1 have been identified in the United States^3^, while Chile has reported a human infection linked to pinnipeds. The global response to the emergence of 2.3.4.4b has identified exposure or infection in a wide variety of mammals including skunks, raccoons, bears, foxes, horses, and mammals^4,5^. Although most spillover infections to mammals are presumed to arise from birds, the species origins, exposure routes and extent of onward transmission in mammals are speculative, frustrating efforts to prevent re-emergence or understand how detections translate to zoonotic risk. Furthermore, most mammals reported infected with 2.3.4.4b are not reservoirs of non-H5 influenza A viruses (IAV)^6^, reducing opportunities for co-infection and reassortment which could alter viral properties and exacerbate pandemic risk. Indeed, most IAV pandemics to date have arisen through inter-species transmission and reassortment, including the 2009 swine-origin H1N1 pandemic^7^.

Bats are reservoirs of prominent viruses with zoonotic potential, such as Marburg virus, Nipah virus, Severe Acute Respiratory Syndrome Coronaviruses, and a variety of rabies-related lyssaviruses^8^. More recent discoveries of bat IAVs, including H9N2 in African bats^9,10^ and H17N10^11^, H18N11^12^ and H18N12^13^ in Central and South American bats, have raised concerns about the zoonotic potential of these viruses. To date, no bat H5N1 infections have been reported, potentially reflecting ecological barriers to exposure between bats and current avian and mammalian reservoirs, physiological barriers to infection in bats such as the distribution of sialic acid receptor types or innate immunological barriers, or limited surveillance effort.

Here we combine an 8-year, multi-site longitudinal study of wild vampire bats, analyses of sialic acid receptor distributions and H5 haemagglutinin binding across bat tissues, and *in vitro* experiments to assess ecological and virological factors shaping bat susceptibility to infection with H5N1 IAVs. We focus on common vampire bats (*Desmodus rotundus*) which, via nightly blood feeding, directly interface with both domestic livestock, marine birds and mammals, potentially including species that experienced mass mortality from 2.3.4.4b^14–17^. We found evidence of H5 IAV infection in vampire bats and identified the likely host species that acted as a source of such exposures. We also proposed feasible pathways for future IAV spread across bat populations and from marine environments to domestic animal species. Finally, we evaluated cellular and molecular determinants relevant to virus transmission or co-infection.

### Recurrent H5 exposure in coastal vampire bats

We first evaluated evidence for H5 IAV exposure in vampire bats by surveying eight coastal (N=804 individual bats) and four Andean sites (N=57) between 2011 and 2024 in Peru (Fig.1a; Extended Data Table 1). Our coastal sites included four putatively marine-associated vampire bat colonies which were located <100m away from the coastline, providing potentially direct access to clade 2.3.4.4b-affected marine mammal and bird populations (high-risk colonies), alongside inland colonies that were located >5 km from the coastline and therefore not expected to be exposed to marine wildlife (moderate-risk colonies). Evidence of IAV infection in inland colonies might therefore signal virus spread within coastal bat populations from the marine-associated colonies. The Andean vampire bat populations included are genetically isolated from coastal populations and thus had no plausible connection to marine IAV hosts or to coastal bats (low-risk colonies)^18^. Our sampling included both pre- and post-epizootic periods (Fig. 1b). Unsurprisingly, no IAV was detected by qPCR in oropharyngeal and rectal swabs from 117 bats captured approximately 8 months after the peak of the epizootic in marine wildlife (October/November 2023). However, anti-H5 antibodies were detected by ELISA in 14 out of 861 serum samples and, strikingly, were found exclusively in marine-associated sites (LMA4, ICA1, ICA2). Seroprevalence in marine sites varied annually from 0-8% (mean: 3.2%, N=431) and exceeded 0% in all sites in 2023 and 2024 (i.e. after the 2022/2023 epizootic). The fourth marine-associated site (LMA10) was abandoned by bats prior to the 2.3.4.4b epizootic, precluding post-epizootic sampling. Pre-epizootic seropositivity in one marine-associated site (LMA4, years 2011*, 2012 and 2015), presumably attributable to non-2.3.4.4b H5 IAVs, indicated virus exchange at the bat-marine wildlife interface was not a peculiarity of the 2023-2024 epizootic. (*borderline positive)

**Fig. 1.**
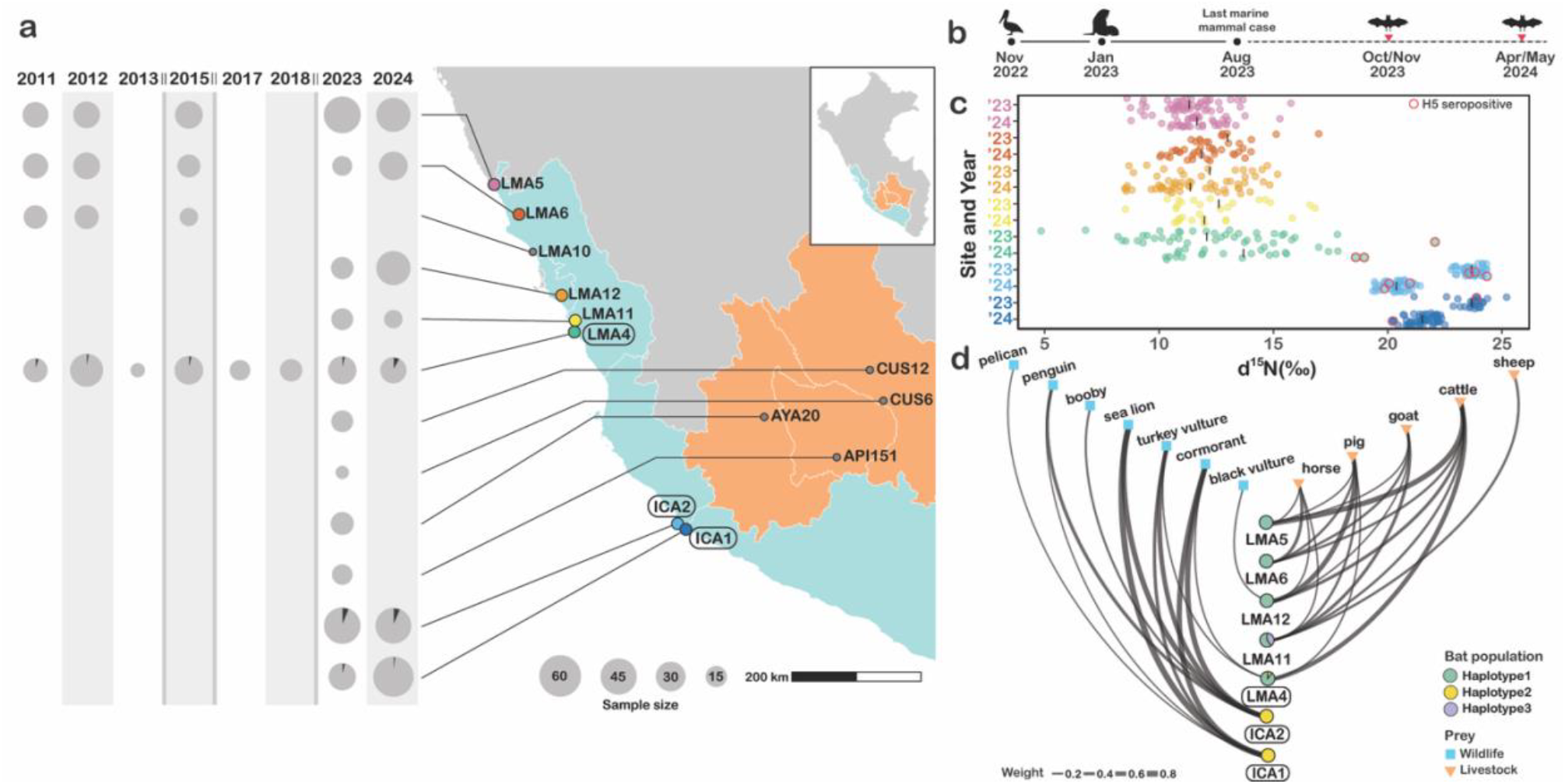
Recurring H5 IAV spillover to marine-feeding vampire bats. **a**, Sampling sites of common vampire bats in coastal (blue) and Andean (orange) regions of Peru. The pie charts indicate H5 seroprevalence in vampire bats between 2011 and 2024. Double bars between years indicate annual sampling gaps. **b**, Timeline of the 2022/2023 epizootic showing first reported infections in birds (November 2022) and South American sea lions (January 2023) and the last reported IAV-infected case in marine mammals (August 2023)^20^. Bat silhouettes indicate post-epizootic bat capture dates. **c**, δ^15^N isotope values across seven sites in 2023 and 2024 (coloured as in panel a). The black vertical lines indicate mean δ^15^N values in each site/year. H5 seropositive individuals are circled in red. High δ^15^N values indicate feeding on higher trophic level prey such as marine wildlife. **d**, Feeding patterns (60 of 72 tested individuals with conclusive prey assignments) and bat population genetic structure in each site (ordered as in panel a), inferred from 12S metabarcoding of vampire bat rectal swabs or blood meal samples. Grey lines show feeding histories of individual bats and pie charts show 12S haplotypes of vampire bats at each site.

To rule out the possibility that bats were exposed to non-H5 IAVs which cross-reacted with our ELISA, we tested sera from ferrets that were experimentally infected with a panel of influenza viruses (Extended Data Table 2). The ELISA detected most H5 subtypes but showed no reactivity to the closely related H1 subtype, the more distantly related H3 subtype or to influenza B virus HA, demonstrating specificity within H5 IAVs. Although our panel of ferret sera did not include H18 IAVs, which have been reported in coastal vampire bats^19^, the large evolutionary divergence between these viruses makes cross-reactivity unlikely. If there was cross-reaction between H18 and H5 antibodies, we would expect similar seroprevalence for both viruses. However, the markedly lower H5 seroprevalence does not support the presence of cross-reactivity (H18: 57.3%, H5: ≤6%; N=286). Consequently, seropositivity to both viruses suggests independent exposures to each virus. Indeed, three of the 14 H5-positive bats (2011, 2012 and 2015) were also H18 positive, suggesting an absence of sterilising cross-protective immunity, and possible coinfection. Taken together, consistent, low H5 seropositivity across three marine-associated sites, and the absence of H5 seroconversion elsewhere, suggests recurring spillover from marine wildlife to bats, but negligible onward transmission within bats.

### Vampire bat H5 exposure linked to marine feeding

Spatial patterns of seropositivity provided circumstantial evidence of marine-derived H5 IAV spillover to wild vampire bats. We next sought to define the precise ecological pathways that facilitated exposure by applying δ^13^C and δ^15^ stable isotope analysis to vampire bat hair, which differentiates long-term (ca. 4-6 months) feeding on wild versus domestic herbivores and prey trophic level, respectively ^21,22^. Among the seven bat colonies sampled in 2023 and 2024, δ^15^N values differed significantly after accounting for year of capture, sex and age of bats (GLM: F_6,456_ = 638.17, P < 0.001). Two of the three marine-associated sites (ICA1: 22.9 ‰ & ICA2: 22.6‰) had much higher mean δ^15^N than all other sites (LMA4, LMA5, LMA6, LMA12, LMA11; Fig. 1c; Tukey HSD: 12.0-13.1‰, all P < 0.001), confirming routine feeding on higher-trophic-level prey, likely reflecting consumption of marine species given their proximity to coastal resources^15^. (Extended Data Fig. 1). Bats from the third marine-associated site (LMA4) had lower mean δ^15^N values than ICA1 and ICA2 (mean: 13.1‰), but exhibited extreme heterogeneity, including individuals with high δ^15^N, suggesting specialised feeding on marine predators (Fig. 1c). Interestingly, the three bats with the highest δ^15^N values in LMA4 (the upper 7% of δ^15^N values in this site) were all seropositive to H5 IAVs. Consistent with marine prey as the source of IAV exposure in bats, stable isotope mixing models using both δ^13^C and δ^15^N values across the seven coastal sites showed that H5 seropositive bats consumed a higher proportion of marine diet (48 ± 0.08–84 ± 0.1%) than H5-negative individuals (6 ± 0.04–69 ± 0.14%).

Crucially, no bats with isotopic evidence of livestock diet, including those which roosted alongside seropositive marine-feeding bats in LMA4, were seropositive. This strongly suggests that H5 exposures in bats represented recurrent spillover from contact with marine wildlife or their environment rather than ongoing circulation within bat colonies (Fig. 1c).

We next used 12S mitochondrial DNA metabarcoding^23,24^ to identify the exact species that might have exposed bats to H5 IAVs and to project which non-marine prey species might be at risk of H5 IAVs if it became established in vampire bat populations. Rectal swabs or blood meal samples from 27 marine-associated bats with conclusive prey assignments contained DNA from a wide range of species including South American sea lion (*Otaria byronia*), neotropic cormorant (*Phalacrocorax brasilianus*), Peruvian pelican (*Pelecanus thagus*), banded penguin (*Spheniscus* sp.), booby (*Sula* sp.), turkey vulture (*Cathartes aura*) and black vulture (*Coragyps atratus*)(Fig. 1d). All five seropositive bats with available metabarcoding data fed on wild birds or marine mammals that were either heavily impacted by the H5N1 epizootic or closely related to the affected species, decisively linking bat diet and IAV exposure^14^. In the isotopically heterogeneous (‘mixed diet’) site LMA4, bats fed on both wild birds (vultures and cormorants) and livestock (pigs and cattle), highlighting the vampire bat’s role as an ecological link between IAV susceptible species in marine and terrestrial agricultural ecosystems (Fig. 1d). As expected, bats in inland sites fed on domestic livestock including horses, pigs, sheep, goats, and cattle (Fig. 1d). At the individual bat level, 27 out of 60 rectal swab samples contained DNA from multiple prey species, including mixtures of avian and marine mammal DNA, suggesting that individual bats might enable virus transmission between otherwise disconnected species even without IAV circulation in bats (Extended Data Fig. 2).

Although our field studies suggested that historical IAV spillovers to bats most likely caused dead-end infections, viral evolution in the marine hosts might increase the future likelihood of bat-to-bat transmission. If this was the case, geographic or ecological isolation of marine-associated bats could curtail further spread to livestock-feeding bat populations. We therefore opportunistically mined our metabarcoding data (which also amplified bat mtDNA) to explore genetic connectivity between populations with and without evidence of H5 exposure, noting that mtDNA is a conservative metric of gene flow in vampire bats^18^ (Fig. 1d). Despite clear genetic separation between livestock- and marine-associated sites, both Haplotype 1 (livestock-associated) and Haplotype 2 (marine-associated) individuals cooccurred at the mixed-diet site (LMA4). Curiously, the sole genotyped bat carrying Haplotype 2 in LMA4 was seropositive and (as shown above) fed on wild birds. Given the geographic distance separating ICA2 from LMA4 (344 km) is more than 6 times the maximum recorded dispersal distance for vampire bats (<55km^25^) this finding suggests that despite the availability of livestock prey in LMA4, cross-generational maintenance of this bat’s ancestral wildlife feeding preference promoted IAV exposure.

### Sialic acid receptors and H5 binding in vampire bat tissues

Having mapped the ecological pathways that caused H5 IAV spillover from marine wildlife to bats, we next explored whether bat physiology might preclude productive IAV infections, which would comprise a potent barrier to onward transmission in bats. IAVs use sialylated glycan receptors to attach to host cells, with the type of sialic acid (SA) linkage (α2,3 and α2,6) determining host susceptibility to avian and human IAVs, respectively^26^. We therefore stained formalin-fixed paraffin-embedded vampire bat tissues (trachea, lung, kidney, liver, intestine) with lectins to characterise IAV receptor distributions^27^ and incubated these tissues with purified HA from A/dairy cow/Texas/24-008749-002-v/2024 to assess binding via immunohistochemistry^28^. α2,6-Gal (human) SA receptors were distributed on the cilia of the trachea (Fig. 2k), and supported strong HA binding (Fig. 2p), with no binding detected in other tissues (Fig. 2l-2o). α2,3-Gal (avian) signals were detected in epithelial cells of the trachea, kidney and intestine (Fig. 2f, 2g, 2j) but generally at low levels. No lectin signal was detected in the lung (Fig. 2i). We note that prolonged storage of these tissues in formalin likely reduced lectin staining and HA binding. We therefore consider positive results as informative, but negative results as inconclusive. We are currently repeating these assays on optimally preserved tissues. Nonetheless, results to date suggests that H5N1 (A/dairy cow/Texas/24-008749-002-v/2024) HA is able to attach to the upper respiratory tract of vampire bats, and that H5N1 viruses would likely be able to use α2,6-Gal SA receptors for binding to these tissues.

**Fig. 2.**
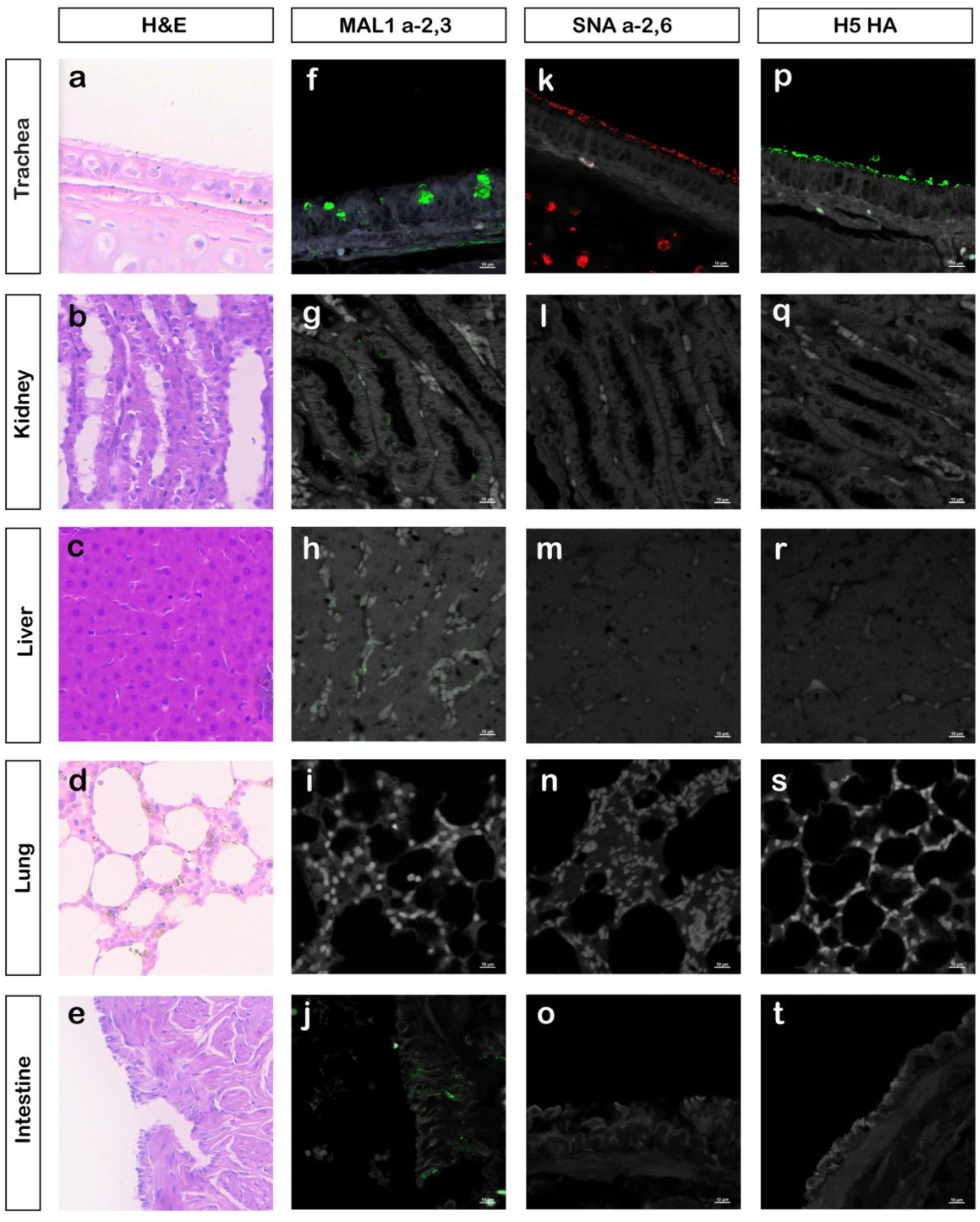
Distribution of avian and human IAV receptors and HA binding in vampire bat tissues. **a-e**, H&E (haematoxylin and eosin) staining of bat tissues. **f-j**, α2,3-SA distribution in each tissue. **k**, α2,6-SA distribution in the cilia and connective tissue of the trachea. **l-o**, absence of α2,6-SA signals in the kidney, liver, lung, and intestine tissues. **p**, H5N1 HA protein binding in the cilia of the trachea. **q-t**, absence of HA signals in the kidney, liver, lung, and intestine tissues. Scale bar: 10µm. MAL, *Maackia amurensis* lectin; SNA, *Sambucus nigra* lectin.

### Vampire bat cells support H5N1 replication

Exposure to a virus and the presence of receptors in physiologically relevant tissues is not sufficient to permit viral infection of cells. To determine if bat cells could support H5N1 IAV replication, we inoculated a panel of vampire bat cell lines (derived from the lung, liver, and kidney) with four H5N1 strains (Extended Data Fig. 3b-3d, 3f-3h; MDCK: Extended Data Fig. 3a & 3e; Fig. 3a & 3b). The strains tested were A/fur_seal/Salisbury_Plain/004762/2024, a mammalian-origin clade 2.3.4.4b; A/brown_skua/Hound_Bay/133947/2023, an avian-origin clade 2.3.4.4b; A/muscovy_duck/England/074477/2021, AIV07, clade 2.3.4.4b from early in the panzootic; and A/Vietnam/1203/2004, a clade 1 virus from a human, preceding the emergence of clade 2.3.4.4b. All four viruses infected (Fig. 3a, 3b, 3d) and replicated (Fig. 3c) in all three bat cell lines, despite the variation in cytopathic effect between strains and in permissivity among cell types. The avian strains generally infected a larger proportion of cells across all three cell lines compared to the mammalian-adapted strain (Fig. 3a). Titration of virus in supernatants by plaque assays showed that all viruses replicated at higher levels in liver cells, particularly the mammalian-adapted strain.

**Fig. 3.**
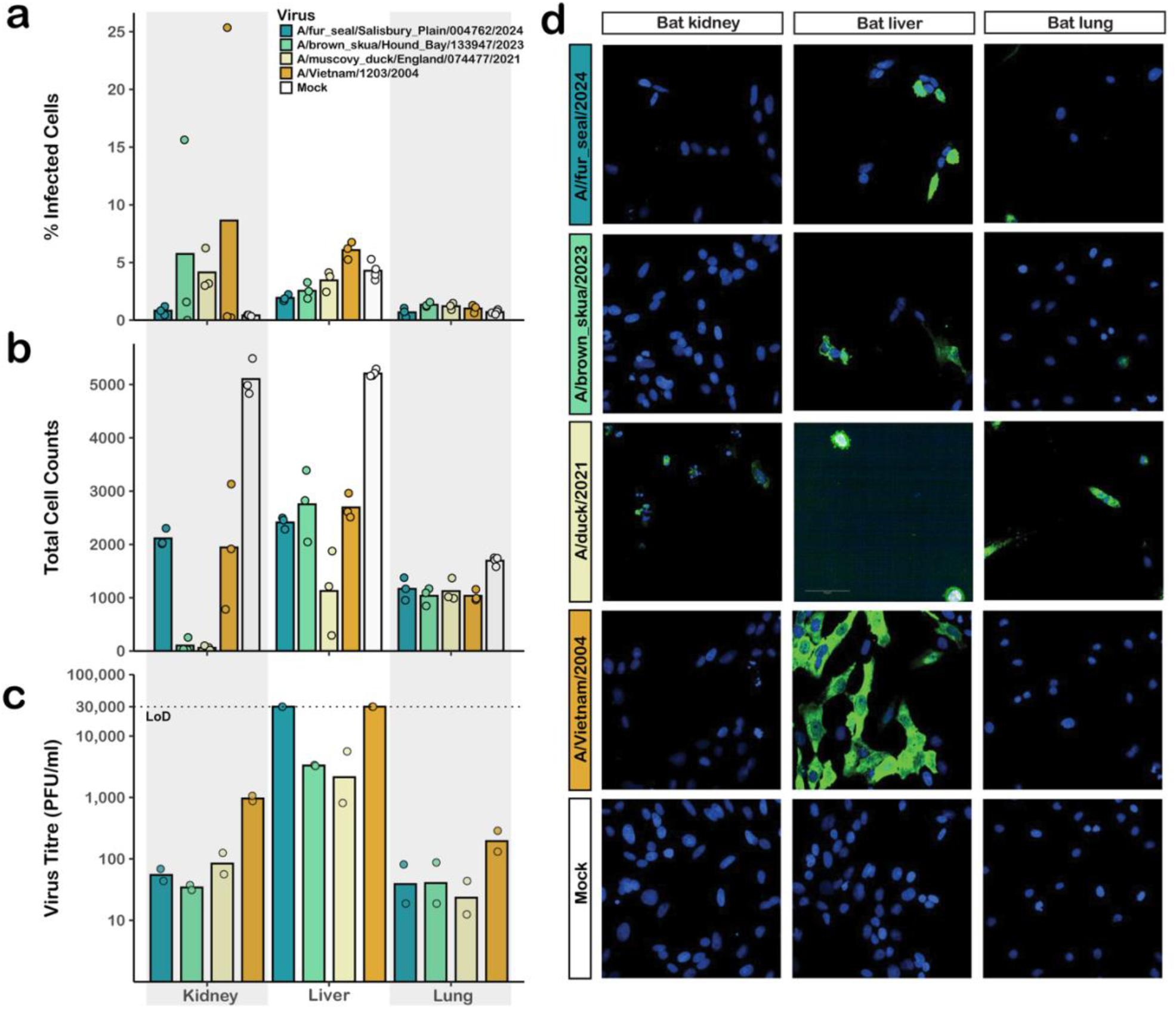
H5N1 IAVs infect and replicate in vampire bat cells. **a**, Percentages of vampire bat cells infected by four H5N1 IAVs at MOI 0.1 PFU/cell. **b**, Total number of DAPI objects (cells), independent of infection. **c**, Plaque titres of supernatants harvested at 20 hours post-infection. The titration was limited to 10^-3^, setting the limit of detection threshold (LoD). **d**, immunofluorescence assay (IFA) confirmed viral infection in bat cell lines.

To further explore the compatibility of IAVs with the intracellular machinery of bat cells vampire bat cells, we used fluorescent-tagged reassortant viruses containing the HA and NA from a laboratory-adapted IAV strain (A/Puerto Rico/8/1934, H1N1) and the internal gene segments of IAVs including A/chicken/England/053052/2021 (AIV07, H5N1), A/dairy cow/Texas/24-008749-001-original/2024 (H5N1, clade 2.3.4.4b), A/Texas/37/2024 (human case, H5N1 clade 2.3.4.4b), and A/mallard/Netherlands/10-Cam/1999 (an H1N1 avian virus passaged in swine cells; Extended Data Table 3). Viral gene expression in all cases demonstrated the capacity of bat cells to permit H1-mediated entry and to support replication driven by internal genes of diverse IAVs. Together, these findings open possibilities for reassortment between H5N1 viruses and other IAVs which may infect the livestock prey of vampire bats (e.g. cattle, pigs) (Supplementary Methods and Extended Data Fig. 4).

## Discussion

By combining ecological field studies, sero-epidemiology, and experimental virology, we demonstrate H5 influenza exposure in bats for the first time and characterise an ecologically and virologically permissive pathway bridging IAV-susceptible marine wildlife, terrestrial wildlife, and livestock. Our data suggest that bat exposures arise through intimate contact with infected marine animals via blood feeding (e.g. exposure to virus in breath, mucus and faeces, and potentially to viraemic blood in cases of systemic infection^29^) or through exposure to contaminated environments while feeding on marine prey. Although seropositivity in marine-feed, but not livestock feeding bats supports recurrent spillover of H5 IAVs rather than bat-to-bat transmission, the presence of avian and mammalian IAV receptors in relevant tissues and the ability of bat cells to support productive infection for a variety of IAVs demonstrates a nascent risk which could be realised by virus evolution or shifts in bat ecology. Establishment of a novel and unusually ecologically connected bat reservoir would promote transmission to existing IAV hosts and heighten pandemic risk.

Our immunohistochemistry and *in vitro* data revealed no obvious barriers to productive infections of bats, raising the question of why H5 viruses appear not to have established in vampire bats, despite repeated opportunities. One possible explanation is that the H5 positive colonies were geographically separated by uninhabitable desert areas, implying that spillover to better connected populations might facilitate spread among bats. However, gene flow among coastal bat populations and the apparent lack of transmission to livestock feeding bats from marine-feeding bats in the same colony suggest factors beyond bat ecology. For example, physiological barriers in bats may allow sufficient replication to induce an antibody response, but insufficient viral shedding to transmit. The dose and route of exposure may also alter infection outcomes. Crucially, the infectivity we observed in our *in vitro* experiments was comparable to that reported for H5N1 viruses in non-adapted host cells (10^2^–10^4^ PFU/ml), despite differences in assay formats (*ex vivo* vs *in vitro*)^30^. Other host defences, such as interferon-mediated innate immunity are likely to provide important barriers to transmission^31,32^. Our study highlights that while vampire bats are ecologically poised to become an IAV reservoir, currently circulating H5N1 IAVs appear unable to fully exploit their cellular machinery or evade their intrinsic and innate immune responses.

Given the dynamic evolution of IAVs, the risk of viral establishment in bats regularly exposed to other infected species should not be ignored. Indeed, H5N1 clade 2.3.4.4b has shown atypical tissue tropism (e.g. mammary tissue in dairy cows) and exceptional capacity to establish in new hosts. Moreover, sustained circulation in marine wildlife could lead to the evolution of IAVs with greater capacity for transmission among bats^31,32^. The interaction of vampire bats with other ecologically relevant IAV hosts (e.g. pigs, cattle, horses) which is well-documented and consistent with our dietary metabarcoding could open the door for co-infection and reassortment events which could rapidly alter the odds of circulation in bats and transmission to other hosts^33–36^. Such co-infection risks appear greatest in seropositive bat populations that feed on a mixture of marine wildlife and terrestrial livestock prey. Gene flow between these and livestock-dependent bat populations provide ecological routes for porcine, bovine or equine IAVs to co-infect bats (Fig. 1a & 1d). Supporting this possibility, our data demonstrated both H5N1 IAVs and H1N1 reassortant viruses replicated in bat cells (Fig. 3 & Extended Data Fig. 4). In addition to conventional IAVs, bat-associated H18 viruses co-circulate within coastal vampire bat populations^19^, including, as demonstrated here, in marine-associated populations (LMA4). Although our data suggests that H5 and H18 IAVs might share a common tissue tropism in the small intestine in vampire bats^37^ (Fig. 2j), differences in receptor usage and packaging signals may make these combinations of viruses unlikely candidates for re-assortment^38,39^. Monitoring bat exposures to other locally relevant IAVs and using an extended panel of viruses would help assess reassortment risks in future work. Finally, the current geographic range of vampire bats stretches into northern Mexico and is predicted to expand into H5N1-affected cattle populations in Texas and New Mexico due to climate change. Assuming H5N1 continues circulating in cattle, the expanding species interface driven by northward bat range expansions or potential southward expansions of H5N1 in cattle might increase opportunities for bovine-associated H5N1 IAVs to emerge in vampire bats ^40–42^.

Several limitations of this study deserve discussion. First, both mammal-adapted (i.e. pinniped) and avian H5 IAVs circulated along the Peruvian coast during our study period^2^, and serological data alone left ambiguity in whether vampire bats were exposed to one or both strains, or low pathogenic H5 strains. As prior mammalian adaptation would be expected to favour transmission among bats, resolving the viral strain(s) involved is important to gauge the risk of future spillover. Circumstantially, H5-exposed bats in marine-feeding sites fed on both marine birds and pinnipeds but H5-exposed bats in the mixed diet site fed on birds only, suggesting direct transmission from birds to bats (Fig 1d; Extended Data Fig. 2). Sequencing of IAV RNA from infected bats would provide decisive strain differentiation but will require future sampling of bats concurrently with epizootics in other species rather than in post-epizootic periods as conducted here. Second, we cannot rule out the possibility that seropositivity in bats reflects repeated antigen exposure rather than genuine infection, though the ecological circumstances of high H5N1 positivity in bat prey, coupled with our laboratory data demonstrating cellular permissiveness suggest the absence of strong barriers to productive infection. Controlled *in vivo* inoculations might provide insights into the infectivity and shedding of H5N1 from vampire bats, however, such experiments cannot be responsibly carried out under currently available biocontainment conditions.

Understanding virus spillover at species interfaces is of paramount importance to pandemic preparedness. Yet, because spillover is shaped by inter-specific interactions that create opportunities for exposure and by host physiological barriers which determine whether exposures lead to infection, assessing spillover risk has been notoriously challenging - requiring multidisciplinary efforts to unify ecological and virological processes^32^. By integrating field surveillance, molecular and chemical ecology, tissue analyses and virological assays, we reconstruct the exposure routes of a potential novel bat host to IAV-affected species, identify onward transmission pathways through high-resolution mapping of species interactions, and experimentally rule out hard physiological barriers to infection. While our current data support limited H5 IAV transmission among bats, the recurrent nature of spillover events, ongoing mammalian adaptation, and prospects for reassortment with other mammalian IAVs, indicate that bats may constitute an open door for future IAV emergence which require closer investigation.

## Methods Experimental design

We tested common vampire bat (*Desmodus rotundus*) sera for H5 IAV across eight coastal sites and four Andean sites in Peru between 2011 and 2024. In addition to sera, oropharyngeal and rectal swabs, hair and bloodmeals were collected from seven out of the eight coastal sites in October/November 2023 and April/May 2024 (LMA10 was not sampled). Six sites located in Lima Department, one in Ica, one at the border of Ica and Arequipa. LMA4, LMA10, ICA1 and ICA2 are less than 100m from the sea front and the rest of the sites are 5km away from the coast. Andean sites consist of two roosts in Cusco and one each in Ayacucho and Apurimac (Extended Data Table 1).

### Field capture

Up to fifty bats were captured at each site over three consecutive nights from 18:00 to 6:00 using mist nets ranging from 3 to 12 meters, depending on the size of the roost entrance. LMA5, LMA6, LMA12, LMA11 are man-made structures (tunnels) whereas LMA4, ICA1 and ICA2 are sea caves. The mist net was monitored every thirty to sixty minutes; the bats were gently removed from the net and kept individually in cotton drawstring bags for a maximum of two hours. When the above methods was not possible (due to low mist net capture), diurnal capture was conducted instead.

### Sampling

Individual body weight was measured using a 100g spring scale. Age (adult, sub-adult, and juvenile), sex, reproductive status (reproductive, non-reproductive, pregnant, lactating), feeding status, and forearm length were recorded before sampling. Oropharyngeal and rectal swabs, blood samples, and hair samples were collected from each bat. The swabs were stored in 2 mL cryovials pre-filled with 1 mL DNA/RNA Shield (Zymo Research, CA, USA) at room temperature for 30 min (for inactivation) before being transferred to dry ice. Blood samples (200 µL) were collected from the cephalic vein using a sodium-heparinised capillary tube after lancing with a 23G needle. The whole blood was kept in a serum separation tube and centrifuged for 3 min using a Starlab Minicentrifuge. The serum samples were then transferred to 0.5 mL microtubes and moved to dry ice. Hair samples (15 mg) were collected from the scapular region of each individual using curved, blunt-end scissors or a paw trimmer.

### Stable isotopes analysis

Hair samples were cleaned using 1 mL of a 2:1 chloroform:methanol solution in an ultrasonic water bath for 30 minutes. The solvent was subsequently removed, and the samples were air-dried. The cleaned samples were then soaked in 1 mL of Milli-Q water, again in an ultrasonic water bath for 30 minutes, water extracted, followed by freeze-drying to remove excess water. Hair samples were weighed to 0.650–0.750 mg and placed individually in tin capsules. Isotopic analysis was performed using a DELTA™ Q Isotope Ratio Mass Spectrometer coupled with the EA IsoLink™ IRMS System (Thermo Fisher Scientific). Mixing models were conducted using *simmr* package^43^ in R studio. Means and SDs of δ^13^C and δ^15^N from livestock prey inputting the models were combined manually rather than using the function in the *simmr* package.

### H5 competitive ELISA

Bat serum samples were tested for H5 exposure using ID Screen® Influenza H5 Antibody Competition 3.0 Multi-species kits (Innovative Diagnostics, France), according to the manufacturer’s protocol. Briefly, samples and controls were diluted 1:10 using dilution buffer. 50 µl of diluted samples and controls were added to corresponding wells of the ELISA plate and incubated at 37°C for 1 h. Following incubation, the plate was washed 3 times with 1:20 diluted wash buffer. The anti-H5-peroxidase (HRP) conjugate concentrate was diluted 1:10 with the dilution buffer and 100 µl was added to each well of the ELISA plate. The plate was incubated at room temperature for 30 min. Upon completion of the incubation the wash step was repeated to remove excess unbound conjugate and 100ul/well 3,3′,5,5′-tetramethylbenzidine (TMB) substrate was added. After 15 min of incubation in the dark, the colour development was terminated by adding 50 µl of stop solution. The optical density was measured at 450nm in a Multiskan FC plate reader (Thermo Fisher). The assay was considered valid if the negative control mean was greater than 0.700 and the mean positive control was less than 30% of the mean negative control. The competition percentage for each sample was determined by calculating sample absorbance divided by absorbance of negative control mean.

### Rectal and oropharyngeal swab extraction

Swabs preserved in DNA/RNA Shield were extracted using *Quick*-DNA/RNA Viral Kit (Zymo Research, CA, USA) following the manufacturer’s protocol. Briefly, samples were removed from −80°C storage and equilibrated to room temperature for 30 min to 2 h. A 200 µL aliquot of each sample was added to 400 µL Viral DNA/RNA buffer, mixed, and transferred to a spin column for centrifugation at 2 min. The column was then transferred to a new collection tube. Next, 500 µL of Viral Wash Buffer was added to the column, centrifuged for 30 s, and the flow-through discarded; this wash step was repeated. Following this, 500 µL of 95-100% ethanol was added, and the column was centrifuged for 1 min. To elute the nucleic acids, 50 µL of DNase/RNase-free water was added to the column, incubated at room temperature for 5 min, and then centrifuged for 30 s.

### cDNA reverse transcription and qPCR Assay

Total nucleic acids extracted from the swabs were reverse transcribed using the MBTuni-12 primer^44^ and the High-Capacity cDNA Reverse Transcription Kit (Applied Biosystems, MA, USA) according to the manufacturer’s protocol on a SimpliAmp thermal cycler (Thermo Fisher Scientific, MA, USA). Real-time PCR targeting the influenza virus M segment was then performed (forward 5’-AAGACAAGACCAATCCTGTCACCTCT-3’; reverse 5’-TCTACGCTGCAGTCCTCGCT-3’; probe FAM-TCACGCTCACCGTGCCCAGTG-TAMRA) using the Brilliant III Ultra-Fast qPCR Master Mix (Agilent Technologies, CA, USA) on a 7500 Fast Real-Time PCR system (Thermo Fisher Scientific, MA, USA). A plasmid-derived influenza M segment was used as the positive control, prepared in ten-fold dilutions ranging from 1E7 to 10 copies. Amplification conditions were set to 95°C for 5 min, followed by 40 cycles of 95°C for 3 s and 60°C for 30 s.

### DNA Metabarcoding

DNA metabarcoding was carried out on DNA extracted from 70 faecal swab samples and from two blood meal samples stored on FTA cards. The corresponding eight faecal swab negative controls and two negative FTA card extraction controls were included. Vertebrate and bird mitochondrial markers were targeted through PCR amplification. For vertebrates, an approx. 95 bp 12S marker (excluding primers) was amplified using the 12SV05 (forward 5’-TTAGATACCCCACTATGC-3’) and 12SV05 (reverse 5’-TAGAACAGGCTCCTCTAG-’3) primer set^23,45^ (hereafter “vertebrate 12S”). For birds, an approx. 260 bp 12S marker (excluding primers) were amplified using the BirT-F (forward 5’-YGGTAAATCYTGTGCCAGC-3’) and BirT-R (reverse 5’-AAGTCCTTAGAGTTTYAAGCGTT-3’) primer set^24^ (hereafter “bird 12S”). Nucleotide tags were added to the 5’ end of the forward and reverse primer of each of the two primer sets to create two sets of uniquely tagged primers^46^. For each primer set, the 7 nucleotides-long tags had at least three mismatches between them. In addition, tags had 1-2 nucleotides at the 5’ end to increase complexity on the flow cell. The workflow principally followed Bohmann et al. 2018^47^. The resulting sequence data were processed separately for the vertebrate 12S and bird 12S datasets, as outlined in Lynggaard et al. 2022^48^.

### Statistical analyses

A linear model was used to compare δ^15^N between sites while controlling for potential confounders. The model included site, year of capture, and bat sex and age as fixed effects. Post hoc pairwise comparisons (Tukey HSD) were then performed to test all site-to-site differences, with p-values adjusted to account for multiple testing.

### Lectin staining and haemagglutinin binding in vampire bat tissues

The haemagglutinin (HA) protein used was from A/dairy cow/Texas/24-008749-002-v/2024 (Cambridge Bioscience, UK), which is the closest commercially available strain to the South American H5 IAVs. Both experiments followed protocols from previously published studies^28,49^. Briefly, vampire bat tissue sections were deparaffinized after paraffin embedding, rehydrated, and incubated with 1% BSA in PBS for 30 min at room temperature (RT) for both lectin staining and HA binding. For lectin staining, sections were retrieved in 0.01M EDTA for 30 min at 95°C. After cooling to RT, the sections were washed three times with PBS and incubated with 20 μg/mL Fluorescein-conjugated MAL-I or CY5-conjugated SNA (Vector Laboratories) overnight at 4°C. For MAL-II staining, biotinylated-conjugate MAL-II was used for detecting O-linked glycans with SA α2-3 GalNAc. After antigen repair and PBS wash, sections were incubated with 20 μg/mL MAL-II overnight at 4°C. After lectin incubation, sections were treated with streptavidin conjugated with FITC (Invitrogen) and incubated for 1 h at 4°C. Finally, stained with Phalloidin-iFluor 647 Reagent (ab176759, 1:500, Abcam) for 30 min, then mounted with ProLong™ Gold Antifade Mountant (P36935, Invitrogen). Images were taken using confocal microscopy.

For HA binding, purified HA protein was mixed with primary antibody (mouse anti-His-tag, MBL) at a molar ratio of 2:1 and incubated on ice for 20 min. The final concentration of HA protein is 50 μg/mL for tissue staining. The pre-complexed HA was evenly applied to vampire bat tissue sections and incubated overnight at 4°C. The sections were then washed three times for 5 min with PBS and incubated with Alexa Fluor 488 goat anti-mouse IgG (1:1000, Invitrogen) for 1 h at RT. The final staining, imaging processes follow the same protocol as the lectin staining.

### H5N1 virus infection in vampire bat cells

MDCK cells were maintained in Dulbecco’s Modified Eagle Media (DMEM) supplemented with 5% FCS. FLuDeRo, DR Kidney, and DR Liver cells were maintained in DMEM supplemented with 10% FCS and 1X non-essential amino acids (Merck, Germany). Four isolates of A(H5N1) were used: A/Vietnam/1203/2004 (VN1203), A/muscovy_duck/England/074477/2021, A/fur_seal/Salisbury_Plain/004762/2024, and A/brown_skua/Hound_Bay/133947/2023. VN1203 was obtained from the Worldwide Influenza Centre, the Francis Crick Institute. The remaining viruses were kindly shared by the WOAH/FAO International Reference Laboratory for Avian Influenza at the Animal and Plant Health Agency. All viruses were propagated by passage in the allantoic cavity of 10-day old embryonated hens’ eggs for 18-24 h at 37°C, clarified by low-speed centrifugation and stored at −80°C until use.

For infection, viruses were diluted to the required MOI in DMEM containing 25mM HEPES and inoculated for 1 h onto the relevant cell type seeded the day prior. Cells were incubated for 20 h at 37°C, 5% CO_2_ before supernatant was stored or cells were fixed by 4% (v/v) formaldehyde for at least one hour. Viruses were quantified by plaque assay, conducted according to well-established methods^50^. Staining was conducted as previously described^51^. Briefly, cells were washed of fixative, blocked and permeabilised with 3% bovine serum albumin (Merck, Germany) with 0.2% Triton X-100 (Merck, Germany) in PBS. Cells were stained using a biotinylated anti-influenza NP antibody (clone 2-8C) produced in-house with an Alexa-488 conjugated to streptavidin (Invitrogen, USA). DAPI was used to stain cellular DNA. Imaging was conducted at 5X using the Operetta and quantified using the associated Harmony software (Perkin Elmer, USA) and 60X using the Opera Phenix (Perkin Elmer, USA).

### Ethics

The University of Glasgow’s School of Biodiversity, One Health and Veterinary Medicine Ethics Committee approved protocols for the capture and handling of bats (ref EA59/23). Collection and exportation permits were granted from the Peruvian authorities (RD-000032-2025-DGGSPFFS-DGSPFS; 015-2023-SERNANP-RNIPG).

Reassortant viral experiments were approved by the local genetic manipulation safety committee at the University of Glasgow (GM223.25.1), and the Health and Safety Executive of the United Kingdom. Reassortants derived by reverse genetics were made with the external glycoproteins of the attenuated vaccine strain A/Puerto Rico/8/1934 (H1N1)^52^ which is mouse adapted and attenuated in humans^53,54^. The PR8 strain used in this study also possesses a strong receptor preference for avian-type α2-3 linked sialic acid^55^. Work with reassortants used in this study was physically segregated from work with mammalian viruses with glycoproteins different from PR8.

## Data Availability and Code Availability

## Acknowledgements

I.T. was supported by the Wellcome Trust (WELLCOTR/WT_PHD_PROGRAMME 218518/Z/19/Z) and stable isotopes analysis carried out with NEIF grant (2857.1024). D.S. was supported by WT Senior Research Fellowship (217221/Z/19/Z), UK Medical Research Council (MC_UU_00034/3) and NSF/BBSRC Ecology, Evolution of Infectious Diseases Program (DEB 2011069, BB/V003798/1). PRM was supported by the UK Medical Research Council (MC_UU_00034/3) and the NSF/BBSRC Ecology, Evolution of Infectious Diseases Program (BB/V004697/1). E.H., P.R.M., and S.K.W. were supported by the UK Medical Research Council (MRC) and Department for Environment, Food and Rural Affairs (Defra, UK) through the consortium grant FluTrailMap-One Health (MR/Y03368X/1). M.P. and M.L.T. were supported by UK Medical Research Council (MC_UU_00034/3) and the FluTrailmap-One Health consortium funding (BB/Y007298/1). C.F. was supported by NERC (NE/V014730/1) and BBSRC-MRC (BB/Y006879/1). R.R. was supported by NSF/BBSRC Ecology and Evolution of Infectious Diseases Program (DEB 2011069, BB/V003798/1). J.G.P was supported by WT Senior Research Fellowship (217221/Z/19/Z). K.B. was supported by Carlsberg Foundation Semper Ardens Accelerate Fellowship (CF21-0411), C.L. was supported by VILLUM FONDEN research grant (VIL41390). R.H. was supported by Francis Crick Institute, Cancer Research UK (CC1114), UK Medical Research Council (CC1114) and Wellcome Trust (CC1114). L.A. was supported by National Institute of Allergy and Infectious Diseases, National Institutes of Health, Department of Health and Human Services (contract no. 75N93021C00015). We thank Walter Campana Quintanilla for assisting in field captures; Callum Magill for sharing influenza qPCR protocol and Alice Broos for laboratory support; Rona McGill for stable isotope technical support; Tina Brand, Pernille Selmer Olsen and Lasse Vinner for metabarcoding sequencing support and infrastructure; Clive Barwick and Kirsty Lawman for providing captive sea lion swabs for early trials; Tom Peacock for discussions of influenza strain selection, and collaborators at APHA and ID.Vet.

## Author contributions

I.T. conceptualised and managed the project, acquired funding, performed the investigation, including data collection, sample extractions, qPCR, stable isotope sample preparation, formal analyses of isotopic and metabarcoding data, visualisation of data and wrote the manuscript.

C.L. performed metabarcoding analysis.

L.A. performed *in vitro* H5N1 virus infection and analysis.

S.K.W. performed *in vitro* reassortant virus infection and analysis.

H.C. performed tissue staining.

S.R. performed H5 ELISA.

M.L.T. developed reassortant virus panel.

M.E.G. conceptualised the project.

R.R. provided bat samples from the Andes region.

J.G.P. provided bat tissues.

C.T. and C.Z. performed data collection and project administration.

W.V.B. and S.C.A. performed project administration.

K.P.R., M.R., I.Q.M., W.C., R.Z.V., J.C.M, J.P.C.N. performed data collection.

A.M.S. and J.F.D. provided bat cells.

G.D. validated the immunohistochemistry results.

C.F. supervised the project, review and edited the manuscript.

E.H. and P.R.M. acquired funding, contributed to the experimental design, and supervised the project.

M.P. acquired funding.

K.B. acquired funding and performed metabarcoding analysis.

R.H. supervised the project.

D.S. conceptualised, supervised the project, acquired funding and led review and editing of the manuscript, assisted by all other authors.

