## Additional information for "Spillover of H5 influenza viruses to vampire bats at the marine-terrestrial interface"

**Additional information**
**Extended data figures**

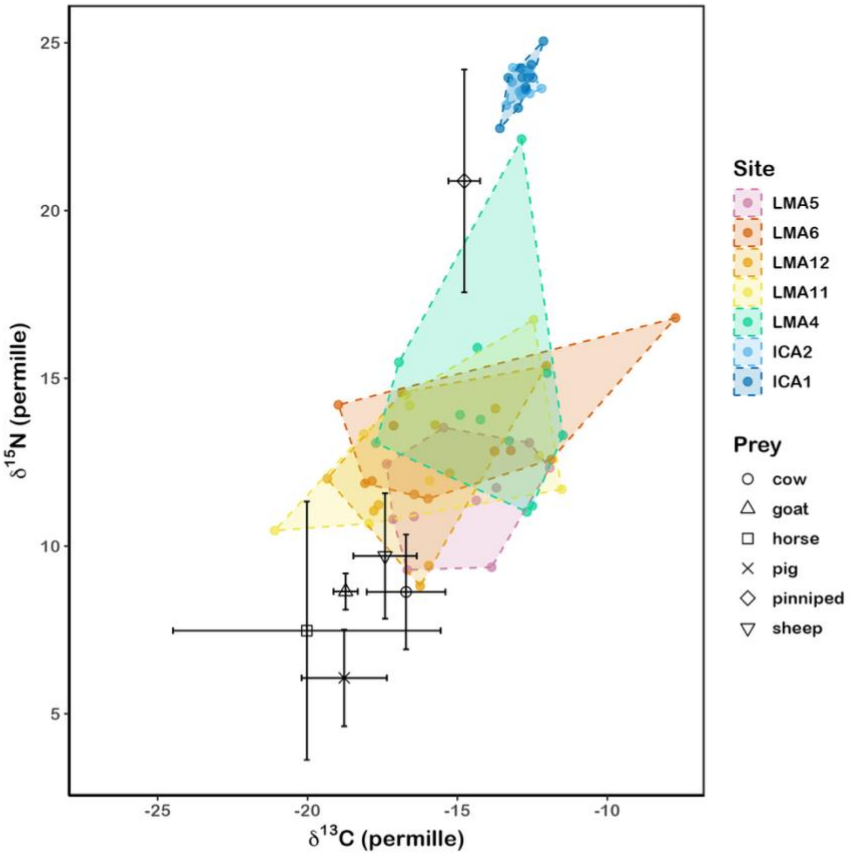


**Extended Data Figure 1 | Distribution of $\delta^{13}C$ and $\delta^{15}N$ isotopic values of 73 bat hair samples collected in 2023.** Each coastal site is colour-coordinated with Fig 1a & 1c. The dashed lines are convex hulls of isotopic space to show variation in diet in each site. Prey isotopic values are presented in different shapes with standard deviation. The isotopic variation in the two marine-associated sites (ICA1 & ICA2) were much lower than the remaining sites. The $\delta^{15}N$ values from these sites were comparable to the pinniped values (diamond-shaped, n=5). While there was a great variation amongst the rest of the sites, the remaining bats had much lower $\delta^{15}N$ which were closer to that of the farm animals.

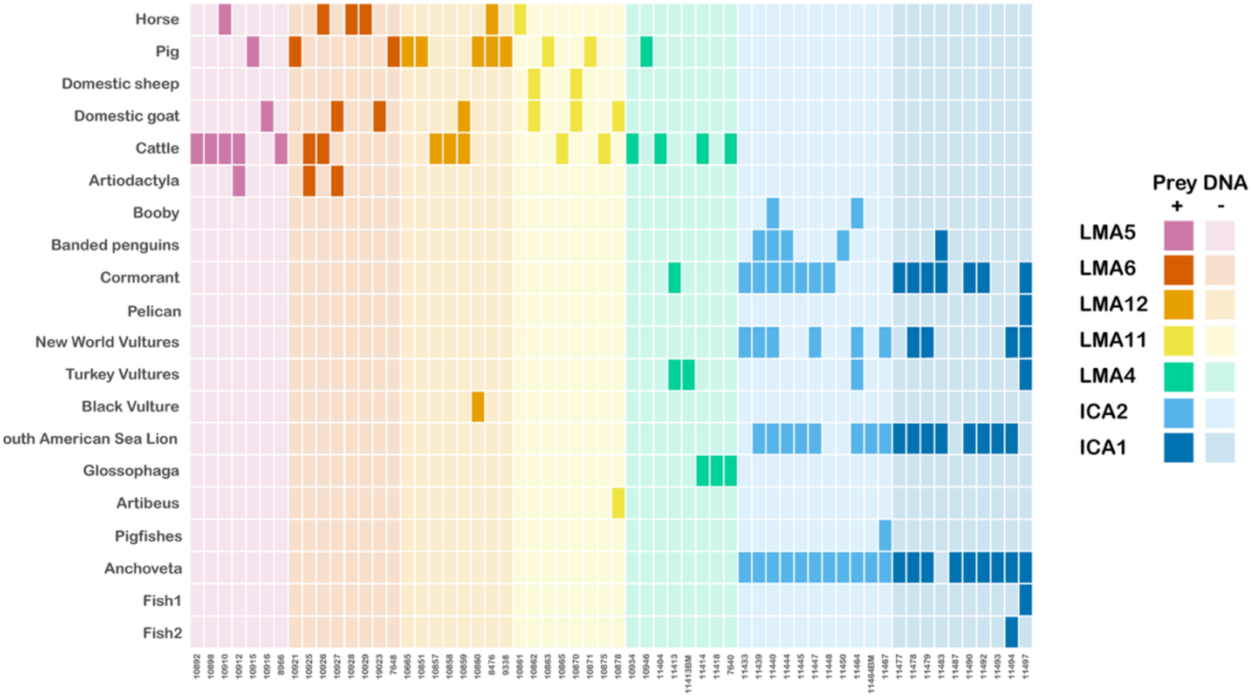


**Extended Data Figure 2 | H5 seropositive vampire bats fed on heavily impacted species during the H5N1 2.3.4.4b epizootic**. We tested 70 individual rectal swabs and two paired bloodmeals from two of the individuals using DNA metabarcoding approach, with an 83% prey DNA detection rate amongst tested samples (60/72). Individual bats are colour coded according to the sites, matching Fig. 1a&1c. Bats from the marine-associated sites (LMA4, ICA1, ICA2) fed on both sea lions and sea birds, including boobies, banded penguins, cormorants, pelicans, turkey vultures and black vultures. These species were hit the hardest during the H5N1 epizootic in Peru. Vampire bats from the remaining sites mainly fed on livestock, which aligned with previous research[1]. The detection of fish and other bat species might be due to environmental contamination. Vampire bats hop on the ground to approach their prey; such behaviour could introduce contamination from prey faeces while swabbing the bats. The two bat species detected shared the same roosts as vampire bats; therefore, environmental contamination is possible.

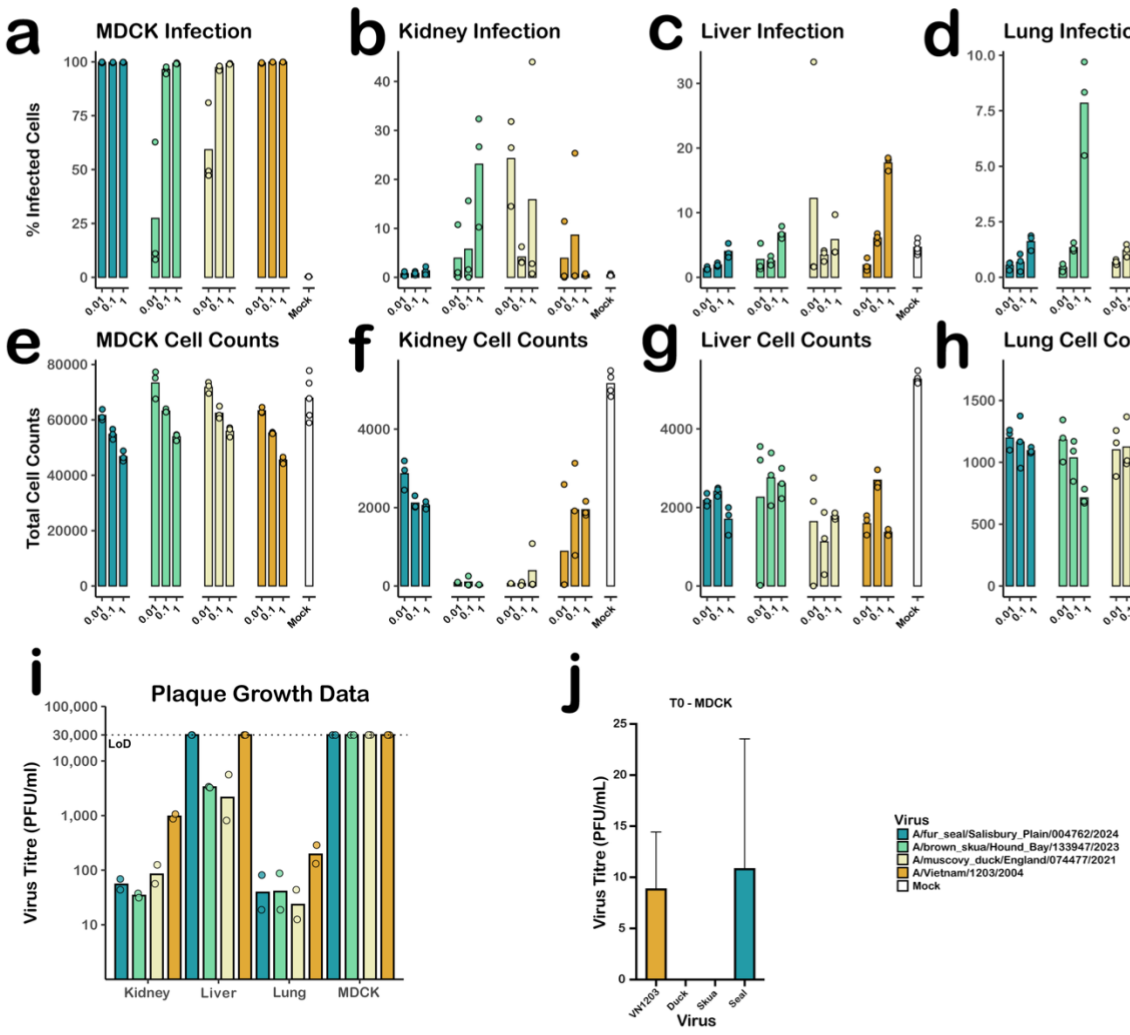


**Extended Data Figure 3 | H5N1 infections in vampire bat cells and MDCK across MOI 0.01, 0.1 and 1.**

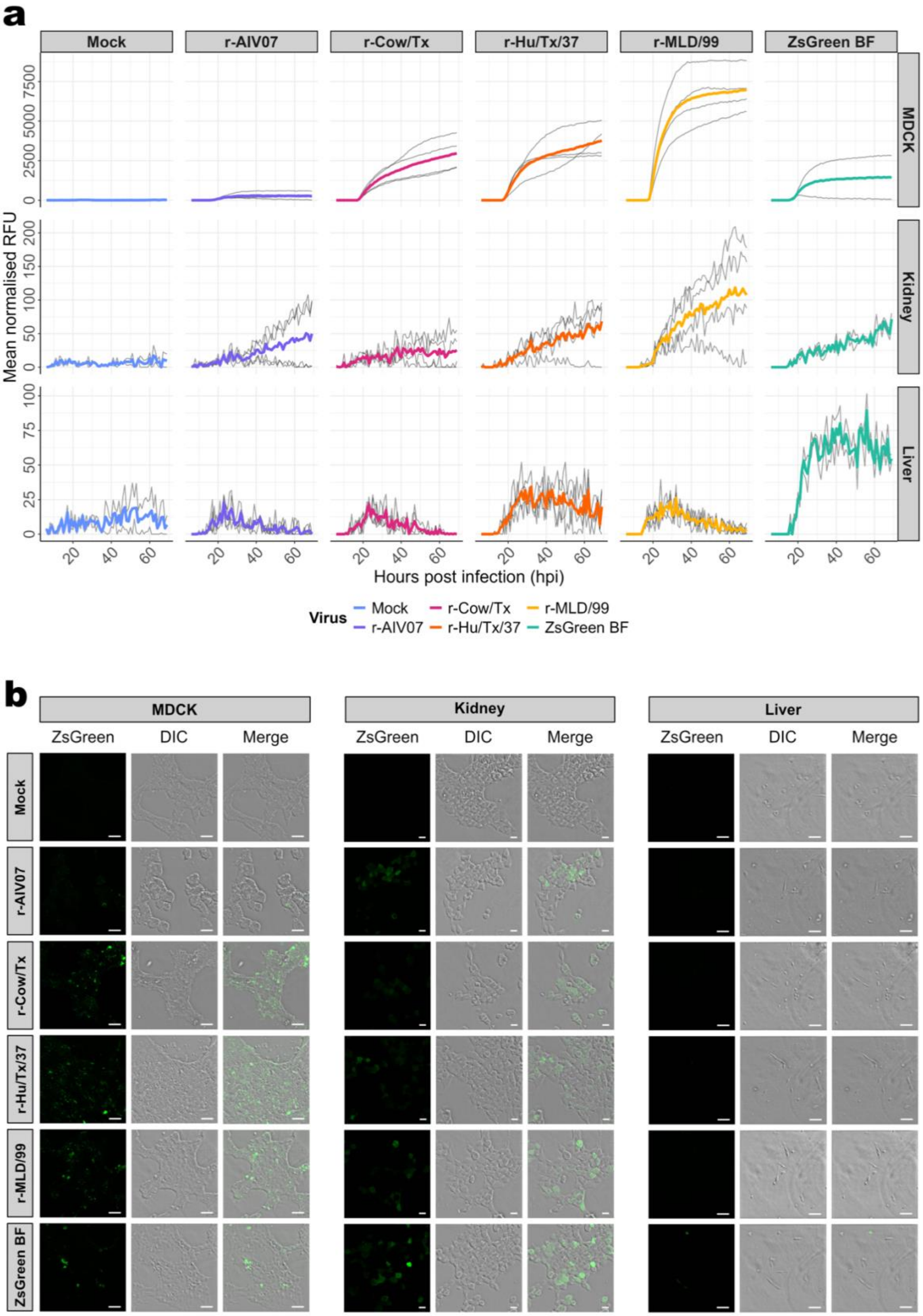
a
Mock
r-AIV07
r-Cow/Tx
r-Hu/Tx/37
r-MLD/99
ZsGreen BF
MDCK
Kidney
Liver
Mean normalised RFU
Hours post infection (hpi)
Virus
Mock
r-AIV07
r-Cow/Tx
r-Hu/Tx/37
r-MLD/99
ZsGreen BF
b
MDCK
Kidney
Liver
ZsGreen
DIC
Merge
Mock
r-AIV07
r-Cow/Tx
r-Hu/Tx/37
r-MLD/99
ZsGreen BF

**Extended Data Figure 4 | Replication kinetics of influenza viruses in *D. rotundus cells***. To investigate the role of H5N1 internal genes in viral replication in *D. rotundus* cells, 6:2 reassortant viruses were generated encoding the HA and NA of the IAV laboratory strain A/Puerto Rico/8/1934 (H1N1) with the remaining genes from a H5N1 or H1N1 virus. Segment 8 of each virus was edited to encode the fluorescent reporter ZsGreen. The viruses were used to infect MDCK cells (as a positive control) and *D. rotundus* cells derived from the kidney and liver. **a**, Cells were infected at an MOI of 0.1 PFU/cell, and replication kinetics assessed over a 66-hour period by measuring reporter gene expression, as normalized Relative Fluorescence Units (RFU). Grey lines indicate independent replicates, with mean fluorescence overlaid in colour. **b**, Confocal micrographs showing ZsGreen reporter gene expression in infected cells, 24-hours post infection at an MOI of 0.1 PFU/ml. Scale bars represent 50 µm for the MDCK and *D. rotundus* liver cells and 20 µm for the D. rotundus kidney cells. For more information, please see Supplementary Materials and Methods.

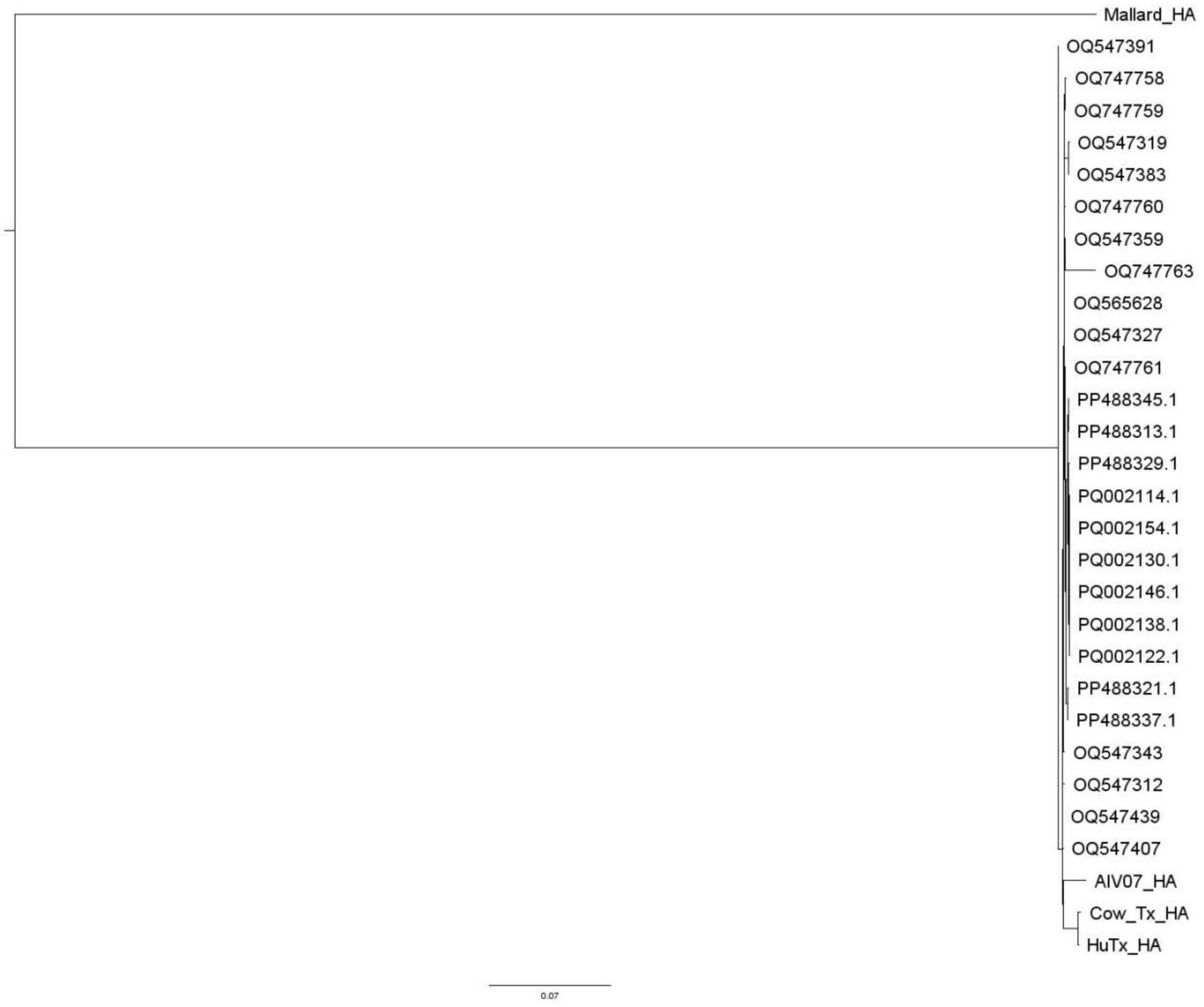


**Extended Data Figure 5 | Maximum-Likelihood phylogeny of viruses used in this study and circulating H5N1 viruses from South America in 2022/2023.** A Maximum-Likelihood phylogeny was constructed to infer the evolutionary relationships between viruses used in this study and H5N1 viruses circulating in Peruvian birds in 2022 and Argentinian seals in 2023.

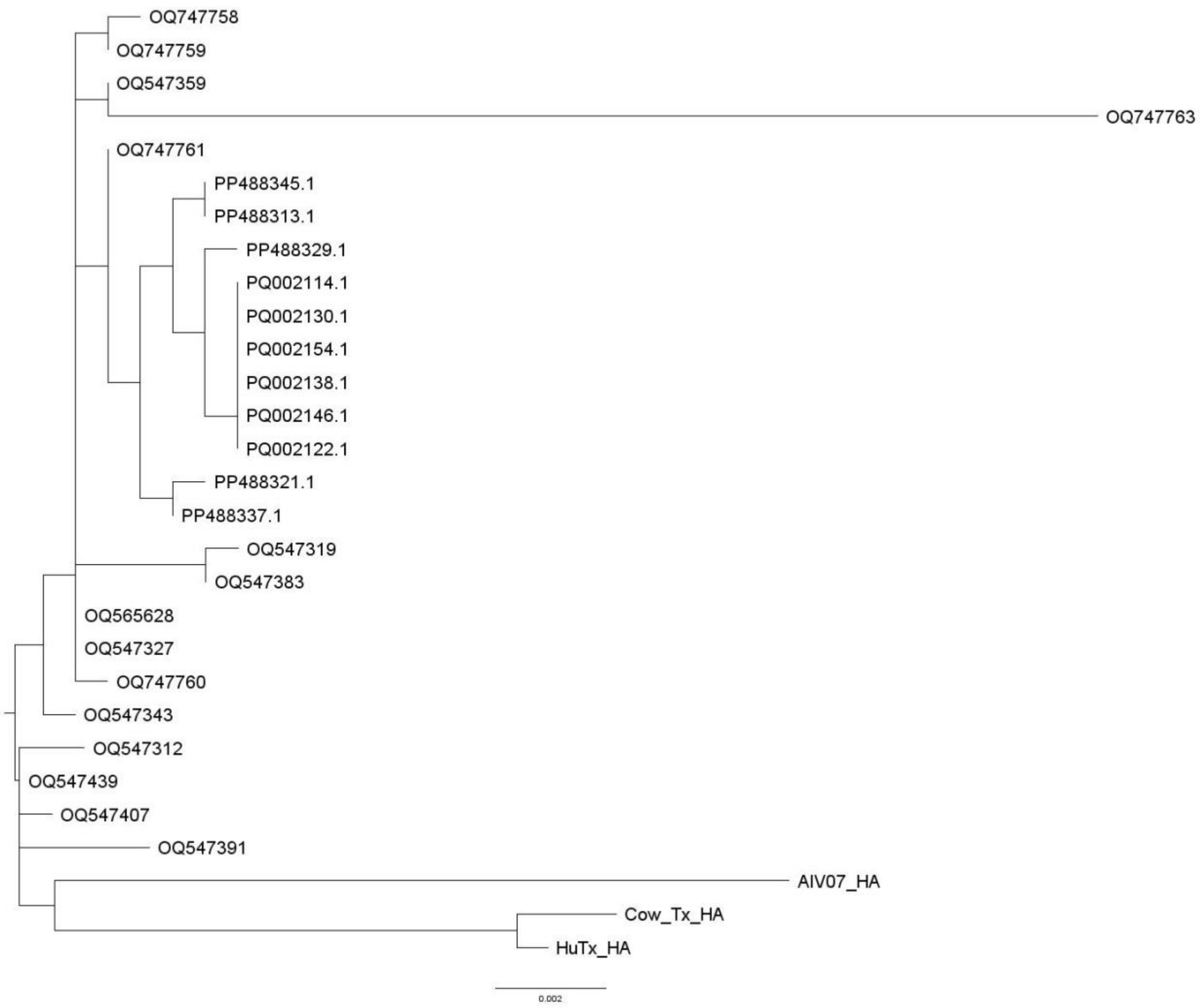


**Extended Data Figure 6 | Maximum-Likelihood phylogeny of viruses used in this study and circulating H5N1 viruses from Peru and Argentina in 2022/2023, without MLD/99.** A Maximum-Likelihood phylogeny was constructed to infer the evolutionary relationships between viruses used in this study and H5N1 viruses circulating in Peruvian birds in 2022 and Argentinian seals in 2023. MLD/99 was excluded from this tree to allow a more precise inference of the relationships between H5N1 strains.

**Extended Data Table 1 | Sites, coordinates, sample types and number of each sample types included in this study.**

| Site | Latitude | Longitude | Sample type | Sampling year | Sample size |
|---|---|---|---|---|---|
| **LMA5** | -10.64 | -77.82 | sera | 2011 | 24 |
| | | | | 2012 | 26 |
| | | | | 2015 | 28 |
| | | | | 2023 | 50 |
| | | | | 2024 | 43 |
| | | | hair | 2023 | 48 |
| | | | | 2024 | 41 |
| **LMA6** | -11.06 | -77.46 | sera | 2011 | 23 |
| | | | | 2012 | 26 |
| | | | | 2015 | 19 |
| | | | | 2023 | 14 |
| | | | | 2024 | 30 |
| | | | hair | 2023 | 15 |
| | | | | 2024 | 30 |
| **LMA10** | -11.59 | -77.28 | sera | 2011 | 20 |
| | | | | 2012 | 20 |
| | | | | 2015 | 12 |
| **LMA12** | -12.18 | -76.85 | sera | 2023 | 18 |
| | | | | 2024 | 43 |
| | | | hair | 2023 | 17 |
| | | | | 2024 | 45 |
| **LMA11** | -12.59 | -76.63 | sera | 2023 | 17 |
| | | | | 2024 | 12 |
| | | | hair | 2023 | 17 |
| | | | | 2024 | 12 |
| **LMA4** | -12.67 | -76.67 | sera | 2011 | 21 |
| | | | | 2012 | 40 |
| | | | | 2013 | 7 |
| | | | | 2015 | 30 |
| | | | | 2017 | 15 |
| | | | | 2018 | 18 |
| | | | | 2023 | 39 |
| | | | | 2024 | 25 |
| | | | hair | 2023 | 39 |
| | | | | 2024 | 25 |
| | | | oral swabs | 2023 | 40 |
| | | | rectal swabs | 2023 | 39 |
| **ICA2** | -15.36 | -75.19 | sera | 2023 | 49 |
| | | | | 2024 | 47 |
| | | | hair | 2023 | 50 |
| | | | | 2024 | 44 |
| | | | oral swabs | 2023 | 50 |
| | | | rectal swabs | 2023 | 50 |
| **ICA1** | -15.44 | -75.07 | sera | 2023 | 27 |
| | | | | 2024 | 60 |
| | | | hair | 2023 | 27 |

| | | | | | |
|---|---|---|---|---|---|
| | | | | 2024 | 57 |
| | | | oral swabs | 2023 | 26 |
| | | | rectal swabs | 2023 | 27 |
| **CUS12** | -13.22 | -72.44 | sera | 2023 | 17 |
| **CUS6** | -13.65 | -72.25 | sera | 2023 | 6 |
| **AYA20** | -13.88 | -73.95 | sera | 2023 | 19 |
| **API151** | -14.44 | -72.91 | sera | 2023 | 15 |

**Extended Data Table 2 | Influenza viruses used to infect ferret, with the ferret sera confirming H5 ELISA specificity.**

| Virus | Type | Strain | IDVet H5 ELISA | Note |
|---|---|---|---|---|
| A/Netherlands/10563/2023 | Influenza A | H1N1 or H3N2 | - | |
| A/Croatia/10136RV/2023 | | H1N1 or H3N2 | - | |
| A/Sydney/856/2023 | | H3N2 | - | |
| A/Cambodia/e0826360/20 | | H3N2 | - | |
| A/California/122/2022 | | H1N1 or H3N2 | - | |
| A/Victoria/4897/2022 | | H3N2 | - | |
| A/Sydney/5/2021 | | H3N2 | - | |
| A/Guangdong-Maonan/SWL1536/2019 | | H1N1pdm09 | - | |
| A/Michigan/45/2015 | | H1N1 | - | |
| A/Norway/31694/2022 | | H1N1 or H3N2 | - | |
| B/Washington/02/2019 | Influenza B | | - | |
| B/Austria/1359417/2021 | | | - | |
| B/Netherlands/10335/2023 | | | - | |
| B/Colorado/06/2017 | | | - | |
| B/Brisbane/60/2008 | | | - | |
| RG71A (A/Astrakhan/3212/2020) | Influenza A | H5N8 | + | |
| A/mink/Spain/22VIR10586-9_3691-3/2022 | Influenza A | H5N1 | - | Consistent with the haemagglutination inhibition test conducted by the Worldwide Influenza Centre, The Francis Crick Institute. |
| A/Cambodia/NPH230032/2023 | Influenza A | H5N1 | + | |

Each sample were tested in triplicates. Our results were consistent with those at the Worldwide Influenza Centre, The Francis Crick Institute.

**Extended Data Table 3 |: Wild-type and reassortant influenza A viruses used in this study.**

| Virus | Origin of external genes | Origin of internal genes | Internal gene NCBI or GISAID reference | Fluorophore | Reference |
|---|---|---|---|---|---|
| PR8 | A/Puerto Rico/8/1934 (H1N1) | A/Puerto Rico/8/1934 (H1N1) | GenBank EF467817 to EF467824 | NA | [2] |
| ZsGreen Bright Flu | A/Puerto Rico/8/1934 (H1N1) | A/Puerto Rico/8/1934 (H1N1) | N/A | ZsGreen inserted between NS1 and NEP flanked by 2A sites. | [3] |
| r-Cow/Tx | A/Puerto Rico/8/1934 (H1N1) | A/dairy cow/Texas/24-008749-001-original/2024 (H5N1) | GISAID EPI_ISL_19014384 | ZsGreen inserted between NS1 and NEP flanked by 2A sites. | This study |
| r-Hu/Tx | A/Puerto Rico/8/1934 (H1N1) | Human A/Texas/37/2024 (H5N1) | GISAID EPI_ISL_19027114 | ZsGreen inserted between NS1 and NEP flanked by 2A sites. | This study |
| r-AIV07 | A/Puerto Rico/8/1934 (H1N1) | A/chicken/England/053052/2021 (H5N1) | GISAID EPI_ISL_9012457 | ZsGreen inserted between NS1 and NEP flanked by 2A sites. | This study |
| r-MLD/99 | A/Puerto Rico/8/1934 (H1N1) | A/mallard/Netherlands/10-Cam/1999 (H1N1) | GenBank KC209512 to KC209519 | ZsGreen inserted between NS1 and NEP flanked by 2A sites. | This study |

**Extended Data Table 4 | GenBank IDs for viruses used in the bioinformatics analysis.**

| GenBank/ GISAID ID | Virus |
|---|---|
| OQ547312 | A/Gallus gallus/Peru/AIS0545/2022 2022/12/03 4 (HA) |
| OQ547319 | A/Gallus gallus/Peru/AIS0547/2022 2022/12/28 4 (HA) |
| OQ547327 | A/Nannopterum brasilianus/Peru/AISA0451/2022 2022/11/22 4 (HA) |
| OQ547343 | A/Gallus gallus/Peru/AIS0539/2022 2022/11/18 4 (HA) |
| OQ547359 | A/Gallus gallus/Peru/AIS0542/2022 2022/12/01 4 (HA) |
| OQ547383 | A/Gallus gallus/Peru/AIS0546/2022 2022/12/18 4 (HA) |
| OQ547391 | A/Gallus gallus/Peru/AIS0548/2022 2022/12/22 4 (HA) |
| OQ547407 | A/Gallus gallus/Peru/AIS0550/2022 2022/12/12 4 (HA) |
| OQ547439 | A/Pelecanus thagus/Peru/AIS0538/2022 2022/11/10 4 (HA) |
| OQ565628 | A/Pelecanus/Peru/VFAR-140/2022 2022/12/4 (HA) |
| OQ747758 | A/Peruvian_pelican/Peru/A074/2022 2022/11/4 (HA) |
| OQ747759 | A/Belcher's_gull/Peru/A102/2022 2022/11/4 (HA) |
| OQ747760 | A/Peruvian_pelican/Peru/A106/2022 2022/11/4 (HA) |
| OQ747761 | A/Belcher's_gull/Peru/A267/2022 2022/12/4 (HA) |
| OQ747763 | A/Western barn owl/Peru/A293/2022 2022/12/4 (HA) |
| PP488313.1 | A/southern elephant seal/Argentina/CH-PD035/2023 (H5N1) (HA) |
| PP488321.1 | A/southern elephant seal/Argentina/CH-PD053/2023 (H5N1) (HA) |
| PP488329.1 | A/South American tern/Argentina/CH-PD030/2023 (H5N1) (HA) |
| PP488337.1 | A/royal tern/Argentina/CH-PD036/2023 (H5N1) (HA) |
| PP488345.1 | A/South American Tern/Argentina/CH-PD037/2023 (H5N1) (HA) |
| PQ002114.1 | A/Southern elephant seal/Peninsula Valades/HA_CH-PD027/2023 (H5N1) (HA) |
| PQ002122.1 | A/Southern elephant seal/Peninsula Valades/HA_CH-PD032_oral/2023 (H5N1) (HA) |
| PQ002130.1 | A/Southern elephant seal/Peninsula Valades/HA_CH-PD032_tracheal/2023 (H5N1) (HA) |
| PQ002138.1 | A/Southern eleyphant seal/Peninsula Valades/HA_CH-PD032_lung/2023 (H5N1) (HA) |
| PQ002146.1 | A/Southern elephant seal/Peninsula Valades/HA_CH-PD032_brain/2023 (H5N1) (HA) |
| PQ002154.1 | A/Southern elephant seal/Peninsula Valades/HA_CH-PD032_rectal/2023 (H5N1) (HA) |
| EPI_ISL_142016 | A/mallard/Netherlands/10-Cam/1999 “MLD/99” |
| EPI_ISL_9012457 | A/chicken/England/053052/2021 (H5N1) “AIV07” |
| EPI_ISL_19027114 | A/Texas/37/2024 (H5N1) |

| | |
|---|---|
| EPI_ISL_19014384 | A/dairy cow/Texas/24-008749-001/2024 (H5N1) |

**Extended Data Table 5 | Significance of the difference in fluorescence over time of different IAVs compared to the negative**.

| Virus | Hours post infection (hpi) | MDCK cells | *D. rotundus kidney* | *D. rotundus liver* |
|---|---|---|---|---|
| Negative | 24 | 0.95 | 0.40 | 0.12 |
| | 48 | 0.99 | 0.61 | 0.03 |
| ZsGreen BF | 24 | < 0.01 | 0.09 | < 0.001 |
| | 48 | 0.02 | 0.01 | < 0.001 |
| r-MLD/99 | 24 | < 0.001 | < 0.001 | < 0.001 |
| | 48 | < 0.001 | < 0.001 | 0.22 |
| r-AIV07 | 24 | 0.43 | 0.04 | < 0.001 |
| | 48 | 0.50 | 0.03 | 0.32 |
| r-Cow/Tx | 24 | < 0.001 | 0.01 | < 0.001 |
| | 48 | < 0.001 | 0.04 | 0.06 |
| r-Hu/Tx/37 | 24 | < 0.001 | < 0.001 | < 0.001 |
| | 48 | < 0.001 | < 0.001 | < 0.001 |

The above table shows the significance values taken from linear models looking at the difference in fluorescence over time (gradient) of infected cells compared to the negative control. Viruses were assessed at two timepoints, 24 hpi and 48 hpi, to assess both single- and multi-cycle replication. p-values are reported to 2 decimal places and significant values ($p < 0.05$) are highlighted in blue.

### Supplementary information

#### Metabarcoding

Prior to the metabarcoding, SYBR Green quantitative PCR (qPCR) was carried out to avoid PCR inhibition and excessive PCR cycles during metabarcoding and to screen negative controls for contamination[4]. The qPCR was carried out on a Mx3005 Real-Time PCR System (Agilent) and included 22 DNA extracts screened with vertebrate 12S and bird 12S primers using undiluted and 1:10 dilution. Further, a positive control (DNA from a bird of paradise, family Paradisaeidae), 9 negative extraction controls and negative qPCR controls were included. For both primer sets, the 25 µL reaction volume included 1 µL DNA template, 1 Unit AmpliTaq Gold, 1x Gold PCR Buffer, and 2.5 mM $MgCl_2$ (all from Applied Biosystems); 0.2 mM dNTP Mix (Invitrogen); 0.5 mg/mL Recombinant Albumina (Bio Labs); 0.6 µM each of 5' nucleotide tagged forward and reverse primer; and 1 µL of SYBR Green/ROX solution [one part SYBR Green I nucleic acid gel stain (S7563) (Invitrogen), four parts ROX Reference Dye (12223-012) (Invitrogen) and 2000 parts high-grade DMSO]. To decrease the amplification of human DNA for the vertebrate 12S primer set, 3 µM human blocker was further included (5'–3'TACCCCACTATGCTTAGCCCTAAACCTCAACAGTTAAATC–spacerC3)[5]. For the vertebrate 12S primer set, the cycling parameters were 95°C for 10 min, followed by 45 cycles of 94°C for 30 s, 51°C for 30 s, and 72°C for 60 s, followed by a dissociation curve. For the bird 12S primer set, cycling parameters were 95°C for 10 min, followed by 45 cycles of 95°C for 30 s, 60°C for 30 s, and 72°C for 60 s, followed by a dissociation curve. The qPCR results showed no signs of PCR inhibition and so metabarcoding proceeded without diluting the sample DNA extracts. For the undiluted sample DNA extracts, the qPCR indicated that the 12S vertebrate primer set required 38 PCR cycles for both sample types, while for the bird 12S primer set, the FTA card samples required less PCR cycles than the faecal swab samples, specifically 30 cycles and 38 cycles, respectively. There was no indication of contamination in the negative controls. DNA metabarcoding was carried out accordingly. The workflow followed the qPCR conditions described above, but omitting SYBR Green/ROX and the dissociation curves. PCR amplifications included four uniquely tagged PCR replicates per DNA extract for the vertebrate 12S primer set and three uniquely tagged PCR replicates per sample for the bird 12S primer set. Alongside the sample extracts, a positive control and the 12 negative extraction controls were included. In addition, a negative PCR control was included for approx. every seven sample extracts. Following PCR amplification, the PCR products were visualized on a 2% agarose gel with GelRed against a 50 bp ladder. The negative controls appeared negative apart from one PCR replicate of one negative extraction control that showed a faint band with the vertebrate 12S primer set. The brightness of the bands were used to approximate the quantity of DNA in each PCR product and were used to guide the pooling of PCR products into amplicon pools prior to library build, this followed Bohmann et al. 2018[6]. The PCR replicates from the 12 negative extraction controls and the negative PCR controls, as well as from the positive control were included. Each amplicon pool included one PCR replicate from one primer set, resulting in seven amplicon pools. The purification of the vertebrate 12S pools was done using a 1.6:1 beads to amplicon pool ratio, whereas for the bird 12S pools was 1:1, both eluted in 35 µL EB buffer (QIAGEN). The subsequent library preparation and quantification of libraries were carried out using the PCR-free Tagsteady library build protocol and followed Carøe and Bohmann 2020[7] and Lynggaard et al. 2023[8], with a final bead purification using a 1:1 bead to library ratio and eluted in 30 µL EB buffer. As such, each sample DNA extract was only subjected to one PCR amplification step prior to sequencing. The resulting seven amplicon pool libraries were pooled equimolarly and sequenced 250 bp paired-end on an Illumina NovaSeq 6000 instrument on a lane of a SP flowcell with 10% PhiX spike-in and other libraries containing markers from different marker regions. Sequencing aimed at approx. 100,000 paired reads for each PCR replicate.

**Bioinformatics**

Firstly, using AdapterRemoval v2.3.3[9], Illumina adapters and low-quality bases were removed and paired reads were merged using Min length 100, min alignment length 50, min quality 28 and quality base 33 parameters. Begum[10] was used to sort sequences based on primer and 5' nucleotide tag sequences, allowing 2 mismatches allowed for primer identifications. After this, Begum was used to filter sequences across the three (bird 12S primer set) or four (vertebrate 12S primer set) PCR replicates from each sample and positive and negative controls. To balance error removal with detection of taxa, we only retained sequences that occurred in minimum two of a sample's three or four PCR replicates in 2 sequence copies[11], and with a minimum length of 200 and 90 for the bird 12S and vertebrate 12S primer set, respectively. Sequences were then clustered into operational taxonomic units (OTUs) using SUMACLUST with a similarity score of 97%[12]. For taxonomic identification, the resulting OTU sequences were manually blasted against NCBI GenBank using 'blastn'. Species level identification was assigned if the sequence had the highest query coverage and the lowest e-value and a 99-100% identity match to a single species; if a 98-98.9% match then it was assigned to genus level, family level if 97-97.9%, order level if 96-96.9%, and class level if 95-95.9% identity match. However, for one OTU in the vertebrate 12S dataset, geographical distribution guided the identification of the sequence as it matched 100% to three species, from which two of them are found only in Australia and New Zealand, and one species is found in the study area. Therefore we assigned this sequence to *Otaria byronia*. To further filter the data, all sequences with less than 99% query cover were filtered out. Thereafter, all diet detections that represented less than 1% and 0.5% of the total diet reads were removed from the bird 12S and vertebrate 12S datasets, respectively. Further, all detections that could not be identified at least to taxonomic order level were filtered out. Finally, detections of the same taxon in the same sample were collapsed.

**Metabarcoding results**

After filtering steps, sequencing data was obtained from 72 samples in both datasets. In the bird 12S dataset, a total of 26,108,270 reads were retained from downstream analyses. From these, 19,612,116 reads belonged to *Desmodus rotundus* (*D. rotundus*) and 6,496,154 reads to non-bat OTUs. From these 72 samples, 24 had both *D. rotundus* and non-bat detections, whereas the rest only had OTUs identified as *D. rotundus*. After removing the bat detections, the number of reads ranged between 14 and 1,685,038 per sample. A total of 8 non-bat OTUs were detected spanning the classes Mammalia (with one OTU identified as *Bos taurus*), and the class Aves represented by six orders, six families, six genera and five species. Two OTUs could not be identified to genus level, whereas two could not be identified to species level. In the vertebrate 12S dataset, 18,943,414 reads were retained after filtering from which 14,808,271 belonged to *D. rotundus*. Vertebrate OTUs not identified as *D. rotundus* were detected in 70 samples. Two samples where *D. rotundus* and another bat were detected (*Artibeus* sp. In one sample, and *Glossophaga soricina* in three). The non-bat OTU detections ranged between 1-5 taxa per sample. After removing all bat detections, 19 non-bat taxa were detected: 3 classes -Actinopterygii- 2 classes, 4 families, 3 genera (1could not be IDed), 1 sp (3 could not be IDed). Within the class Aves - 4 orders, 5 families, 6 genera, 1 species (6 could not be IDed). Mammalia - 3 orders, 4 families (2 OTUs could not be ID), 6 genera (2 could not be IDed), 5 species (3 could not be IDed). In both datasets there were no sequences detected in the negative controls, nor the sequences from the positive control were detected elsewhere.

**Generation of 6:2 reassortant viruses and subsequent infections**

**Cells**

Madin-Darby Canine Kidney (MDCK) cells were maintained in Dulbecco's Modified Eagle Medium (DMEM; Gibco) supplemented with 10% fetal bovine serum (FBS; Gibco). *Desmodus rotundus (D. rotundus)* liver and kidney cells (provided by Prof. Jan Felix Drexler at the Charité Universitätsmedizin Berlin) were cultured in DMEM supplemented with 10% FBS and 1% non-essential amino acids (NEAA; Gibco). All cells were maintained at 37°C and 5% $CO_2$ in a humidified environment.

**6:2 reassortant viruses**

ZsGreen Bright Flu was rescued as previously described (5). Briefly, ZsGreen Bright Flu expresses the ZsGreen fluorescent protein under the control of a modified non-structural protein (NS) segment, where the NS1-2A-ZsGreen-2A-NEP cassette replaces the native NS gene. To investigate the permissiveness of *D. rotundus* cells to genes of HPAI-origin, a series of 6:2 reassortants were generated by combining the external genes of A/Puerto Rico/8/1934 (H1N1) with internal genes of one of four H5N1 viruses, as previously described (6). In these reassortants, the NS segments of the H5N1 viruses were similarly modified to express ZsGreen using the same strategy as ZsGreen Bright Flu, replacing the native NS gene with the NS1-2A-ZsGreen-2A-NEP cassette. For a complete list of viruses used in these experiments, please see Extended Data Table 3. All virus stocks were rescued as previously described (6), propagated at low multiplicity of infection (MOI) in virus growth media (VGM) (DMEM with 1 µg/mL TPCK-trypsin), and infectious titers - expressed as plaque forming units (PFU/mL) - determined using conventional plaque assays (7).

To evaluate the evolutionary relationships between our viruses and those circulating in South American birds and mammals in 2022, all available Peruvian H5N1 hemagglutinin (HA) nucleotide sequences from 2020 to 2024 (n = 15) and Argentinian elephant seal H5N1 HA sequences from 2022 (n = 11) were retrieved from NCBI (Extended Data Table 4). Additional HA nucleotide sequences, including A/chicken/England/053052/2021 (H5N1) ["AIV07"], human A/Texas/37/2024 (H5N1) ["Hu/Tx/37"], A/dairy cow/Texas/24-008749-001-original/2024 (H5N1) ["Cow/Tx"], and A/mallard/Netherlands/10-Cam/1999 (H1N1) ["MLD/99"], were obtained from the GISAID EpiFlu Database (Extended Data Table 4). All HA nucleotide sequences were aligned by translation using Clustal-W with default parameters. Maximum Likelihood phylogenies were inferred from the nucleotide alignments using IQ-TREE 2, also with default parameters. Two phylogenies were constructed: one including MLD/99 to assess the broader evolutionary relationships between the H1N1 and H5N1 strains (Extended Data Figure 5), and another excluding MLD/99 to provide a more precise estimation of relationships among H5N1 sequences while minimising long-branch attraction and outgroup effects (Extended Data Figure 6). The overall topology of the two phylogenies was broadly similar, with small differences in the branching of sequences near to MLD/99 but no substantial change to the major groupings.

**Confocal infections**

MDCK cells and *D. rotundus* liver and kidney cells were seeded onto 13mm, Poly-D-Lysine coated glass coverslips (Avantor) in a 24-well plate (Corning) at a density of $1x10^5$ cells/well. The cells were left to incubate for 24 hours before they were washed twice with DPBS and infected with either VGM (negative control) or one of the five viruses at an MOI of 0.1. Cells were returned to the incubator and left for a further 24 hours before being fixed with 4% (v/v) formaldehyde diluted in DPBS for 10 minutes and washed a further three times with DPBS + 2% BSA. Coverslips were mounted on glass slides (VWR) and imaged with a Zeiss LSM 710 confocal microscope on an AxioObserver platform (Plan-Apochromat 40x/1.4 Oil DIC objective). Simultaneous fluorescence and DIC images were acquired using a 405 nm laser at 2% power, with fluorescence emission from ZsGreen collected by a photomultiplier tube (PMT) detector. Image acquisition and processing were conducted using Zeiss ZEN 3.10 software, with uniform exposure, gain, brightness, and contrast for each cell type.

**ClarioSTAR infections**

Nunc MicroWell 96-Well Optical-Bottom Plates (Thermo Scientific) were seeded with $1.2x10^5$ cells per well in Dulbecco's Modified Eagle Medium (DMEM) supplemented with 10% fetal bovine serum (FBS) and incubated overnight at 37°C with 5% $CO_2$. After 24 hours, the DMEM was removed from the cells and replaced with FluroBright (Gibco, supplemented with 2% L-Glutamine [Gibco] and 0.7% bovine serum albumin [Gibco]) and returned to the incubator for 5 minutes to remove as much residual phenol red as possible. Working stocks of each virus were diluted in FluroBright supplemented with 1gu/µL of VGM to a final concentration of $6x10^4$ PFU/mL. The existing media was aspirated from the 96-well and replaced with 200µL of the appropriate virus inoculum, providing a final MOI of 0.1.

The plate was transferred to a ClarioSTAR microplate reader (BMG Labtech) and incubated for approximately 66 hours at 37°C with 5% $CO_2$. ZsGreen expression was monitored by measuring fluorescence using an excitation wavelength of 470 nm (±15 nm bandwidth) and an emission wavelength of 515 nm (±20 nm bandwidth). The dichroic mirror was set to auto mode with a cutoff at 491.2 nm. The focal height was adjusted to 4.4 mm, and measurements were taken every 15 minutes using a 3 × 3 matrix scan per well. Two independent replicates, each involving separate virus dilutions and infections, were performed for each cell line infected with ZsGreen Bright Flu. Four independent replicates were conducted for each cell line infected with the 6:2 viruses.

Fluorescence data was analysed using R v4.1.0. In brief, data were processed to correct for background auto-fluorescence and for any observed increases in auto-fluorescence over time due to cell death. The background auto-fluorescence in each well was corrected for by subtracting the average fluorescence from timepoints 50–100 from all subsequent readings for that well. To adjust for increasing auto-fluorescence over time, a linear model was fitted to the negative control wells of each cell type to estimate the change in fluorescence due to natural cell death, which was then subtracted from all readings for that cell type.

Due to high heterogeneity in the observed shape of the growth curves, fitting a single model to the data was challenging. Alternatively, fluorescence data were analysed using linear models to calculate the gradient of the growth curves over time. The resulting gradients were compared to the negative control to identify viruses that exhibited statistically significant increases in fluorescence. Two key timepoints, 24 and 48 hpi, were assessed. The 24 hpi timepoint was selected to capture fluorescence following a single cycle of infection and ZsGreen expression, while the 48 hpi timepoint provided insight into fluorescence after multi-cycle replication (Extended Data Table 5).
